# Phagocytic Uptake of Particles by Immune cells Under Flow Conditions

**DOI:** 10.1101/2021.04.01.438153

**Authors:** Megha Srinivas, Preeti Sharma, Siddharth Jhunjhunwala

## Abstract

Particles injected intravenously are thought to be cleared by macrophages residing in the liver and spleen, but they also encounter circulating immune cells. It remains to be determined if the circulating cells can take up particles while flowing in the blood. Here, we use an *in vitro* peristaltic pump setup that mimics pulsatile blood flow to establish if immune cells take up particles under constant fluidic flow. Our results show that the immune cells do phagocytose under flow conditions, and the uptake depends on the cell type, particle size, and flow rate. We demonstrate that cell lines representing myeloid cells, and primary neutrophils and monocytes are similar or better at taking up sub-micrometer-sized particles under flow compared to static conditions. Experiments with whole blood show that even under the crowding effects of red blood cells, neutrophils and monocytes take up particles while flowing. These data suggest that therapeutics may be delivered to circulating immune cells using particulate delivery systems.

## Introduction

A small proportion of nano- and sub-micro-sized particles administered through the intravenous route reach their target cell or organ due to clearance by immune cells^1–4^. The removal of particles from circulation is thought to occur primarily through the phagocytic action of macrophages residing in the liver and spleen^1,5^. Enhanced clearance in these organs appears to be due to the presence of large numbers of phagocytic macrophages and lower blood flow velocity (and hence nanoparticle flow velocity) in the liver sinusoids^6^. However, there are several phagocytic immune cells in circulation, and it remains unclear how these cells interact with the intravenously injected particles.

Phagocytosis of particles by immune cells has primarily been studied using models where the cells and particles are static^7–10^, or when the cells are static, and the particles are flowing^11,12^. In comparison, uptake studies when both the cells and the particles are flowing remains poorly explored. Minasyan suggests that phagocytosis in the bloodstream is likely to be impossible due to the high velocity of blood flow, insufficient time for particulate capture and engulfment by immune cells, and crowding due to a large number of red blood cells^13^, though experiments to prove this have not been conducted. Contrastingly, Tsoi et al.^6^ have shown that peripheral blood mononuclear cells do phagocytose nanoparticles under flow conditions *in vivo*, but their phagocytic capacity may be lower than static conditions. Another *in vivo* study in mice showed that neutrophils in circulation could phagocytose intravenously injected particles^14^. Similarly, others have demonstrated that intravenously injected particles do get taken up by immune cells^15–17^. While these reports suggest that phagocytosis occurs under flow conditions, it does not prove it definitively. One possible alternate explanation is that the particles are taken up by stationary immune cells present in tissues (such as the liver or lung), which then enter circulation again^18^. Though cells with particles may be retrieved from the blood, one cannot be sure of the site of phagocytosis.

Herein, we sought to answer whether phagocytosis may occur when both cells and particles are flowing using an *in vitro* setup. We used a peristaltic pump to mimic the pulsatile nature of blood flow. Experiments on phagocytosis under flow conditions were conducted by varying flow rates from 0.1 to 10 ml/min, which are in the range of flow rates observed *in vivo*^19,20^. Our results show that uptake does occur under flow conditions and that the level of uptake depends on cell type, particle size, and flow rate.

## Materials and Methods

### Cell Lines

RAW 264.7 cells (Merck, USA) were cultured in DMEM complete media (DMEM (CellClone, India) containing 10% fetal bovine serum (FBS, Thermofisher, USA) and antibiotics). Cells were passaged when the flasks were about 80% confluent. The HL-60 cells (ATCC, USA) were cultured in IMDM complete media (IMDM (Merck) containing 20% FBS and antibiotics). Cells were passaged when the cells were at a concentration of 1 million cells in 1 ml.

### Human Blood, Neutrophils and PBMCs

Peripheral venous blood samples were obtained from healthy volunteers following the norms laid out by the Indian Institute of Science’s ethics committee for research with human samples (IHEC No: 5-15032017 and 2-31082018) and with informed consent. The blood was collected in EDTA-coated tubes for purified-cell experiments and citrate, or EDTA-coated tubes for whole blood experiments. For isolating immune cells, the blood sample was loaded onto histopaque (density 1.077 g/ml, Merck, USA) gently and was spun down at 400 RCF for 20 min at room temperature without brake. The buffy coat containing the PBMCs was carefully removed and subjected to RBC lysis for 3 minutes. The bottom-most layer containing the RBCs and neutrophils was subjected to RBC lysis for 9 minutes to obtain the neutrophils. The lysis was quenched with over two-fold volume of 1X PBS. Following quenching, the samples were spun down at 400 RCF for 4 minutes at 4°C. The supernatant was discarded, and the cells were suspended in DMEM complete media. The cells were counted using a hemocytometer, and the required number of cells were used for the experiments.

### Particles

Polystyrene particles (Bangs Laboratories, USA) of varying sizes were used as a model particle system. Volumetrically equivalent amounts of different particle sizes were used, which approximately corresponds to one particle (2.9 µm diameter) ≈ two hundred particles (500 nm diameter) ≈ three thousand particles (200 nm diameter)

### Uptake Studies under Static Conditions

The cells were seeded in a 24-well plate at a concentration of 0.1 or 0.2 million cells per well. Adherent cells (RAW and PBMC) were allowed to adhere for a minimum time of 1 hour, while non-adherent cells (HL-60 and neutrophils) were allowed to remain in the plate for 20 min at 37°C. Then the plates were moved to 25°C (room temperature) or stayed at 37°C. Separately, polystyrene particles (varying numbers depending on the particle size) were sonicated for 5 min and then added to the cells. At the specific time points (2 hours or 4 hours), the media was removed, and cells collected from the wells were washed and stained with propidium iodide (2 µg/ml) to distinguish live from dead and run on a flow cytometer (BD FACSCelesta).

### Uptake Studies under Flow Conditions

A Masterflex L/S easy load peristaltic pump (Cole-Parmer, India) was used for the peristaltic flow experiments along with platinum-cured Silicone Tubing (Sani-tech STHT-C-062-2, 1/16” x 3/16”, Saint-Gobain, India). The tubing was passivated with a filter-sterilized 1% bovine serum albumin in PBS solution for 30 min each time. The tubing was sanitized with a 70% ethanol wash and a water wash before and after each run, followed by autoclaving after a maximum of two runs. Cells were suspended at a concentration of 1 million cells/ml in 1.2 ml of the appropriate type of media. Immediately after adding particles to these cells, the cell and particle suspension was subjected to flow in the peristaltic pump at the desired flow rate. At pre-specified times, the cells flowing through the peristaltic pump were collected, washed, and suspended in 500 µl of 1X PBS. A small aliquot was taken for manual counting (using trypan blue to check for survival). The cell suspension was stained with propidium iodide (2 µg/ml) and run on the flow cytometer. To ensure that the particles had been internalized, trypan blue was added to the cell suspension (in addition to propidium iodide) containing buffer before running on the flow cytometer as trypan blue is known to quench the fluorescence signal of particles stuck to the membrane (and internalized particles are protected from the quenching).

### Whole blood experiments

Heparin (20 IU/ml) was added to the blood sample collected in citrate or EDTA coated tubes within 20 min of collection. Whole blood (500 µl) was plated in a 24-well plate and placed in the incubator at 37 °C for 20 min. Polystyrene particles (500 nm) were sonicated for 5 min and then added to the cells at an approximate ratio of 1:200 (cells to particles) according to the combined theoretical neutrophil (5 million/ml) and PBMC (2 million/ml) counts in the blood. Whole blood and particle suspension was subjected to flow in the peristaltic pump at the desired flow rate for 2 hours. The cells were then collected and subjected to RBC lysis for 8 min. The cells were spun down and resuspended in flow cytometry buffer (PBS containing 1% BSA and 4mM EDTA). They were stained with antibodies against CD45 (clone 2D1, BD Biosciences, USA), CD14 (clone M5E2, BD Biosciences) and CD15 (clone HI98, BD Biosciences) for 20 min at 4°C. The cell suspension was stained with propidium iodide (2 µg/ml) and run on the flow cytometer. All flow cytometry data analysis was performed using FlowJo (Treestar, USA).

### Statistics

The analysis of data and plotting of graphs was done using GraphPad Prism 8 (GraphPad Software, USA). Scatter plot data is reported through a single mean value with a standard deviation. For comparison of two groups, paired Student’s ‘t’ test was used (pairing was based on experiment’s date when cells were from the same passage or individual, and particle additions were from the same stock solution). One-way ANOVA followed by post – hoc Tukey test was used to compare multiple groups.

## Results

Blood flows at different rates (and velocity) in the arteries, veins, and capillaries^19^. To mimic flow rates in each of these circulatory system locations, we chose to perform experiments at three different flow rates – 0.1, 1, and 10 ml/min (∼0.83 mm/s, ∼8.3 mm/s, and ∼83 mm/s). Under these conditions, we measured the uptake of polystyrene particles by immune cells and compared it to conditions where both cells and particles are stationary. Uptake was determined as the percentage of cells that have taken up particles (a measure of how many cells are phagocytic) and the number of particles per cell (average values based on intracellular fluorescence intensities). Further, as the peristaltic setup could not be placed inside a 37°C incubator, studies were performed at room temperature. We determined that for the short time-frame studies (∼2 hours), uptake at room temperature (∼25°C) was slightly (and significantly) lower than 37°C for the adherent macrophage (RAW 264.7) cell line and not significantly different for the suspension myeloid (HL-60) cell line (Suppl. Fig. 1).

### Uptake by an adherent cell line

The adherent mouse macrophage cell line, RAW 264.7, is highly phagocytic across particle sizes under static conditions^21^. To determine this cell’s ability to take up particles while flowing, we tested uptake using polystyrene particles of three different diameters – 200 nm, 500 nm, and 2900 nm (Suppl. Fig. 2). We observed that uptake under flow conditions was dependent on both particle size and flow rate (Figure 1). The 200 nm particles were taken up equally well under both static and flow conditions, and no significant differences were observed across flow rates (Figure 1A and 1B). On the other hand, the percentage of cells that took up the 500 nm particles was greater at 1 ml/min flow rate than the static condition and other flow rates (Figure 1C). However, the number of particles per cell was not different across flow rates and compared to the static condition (Figure 1D).

**Figure 1:**
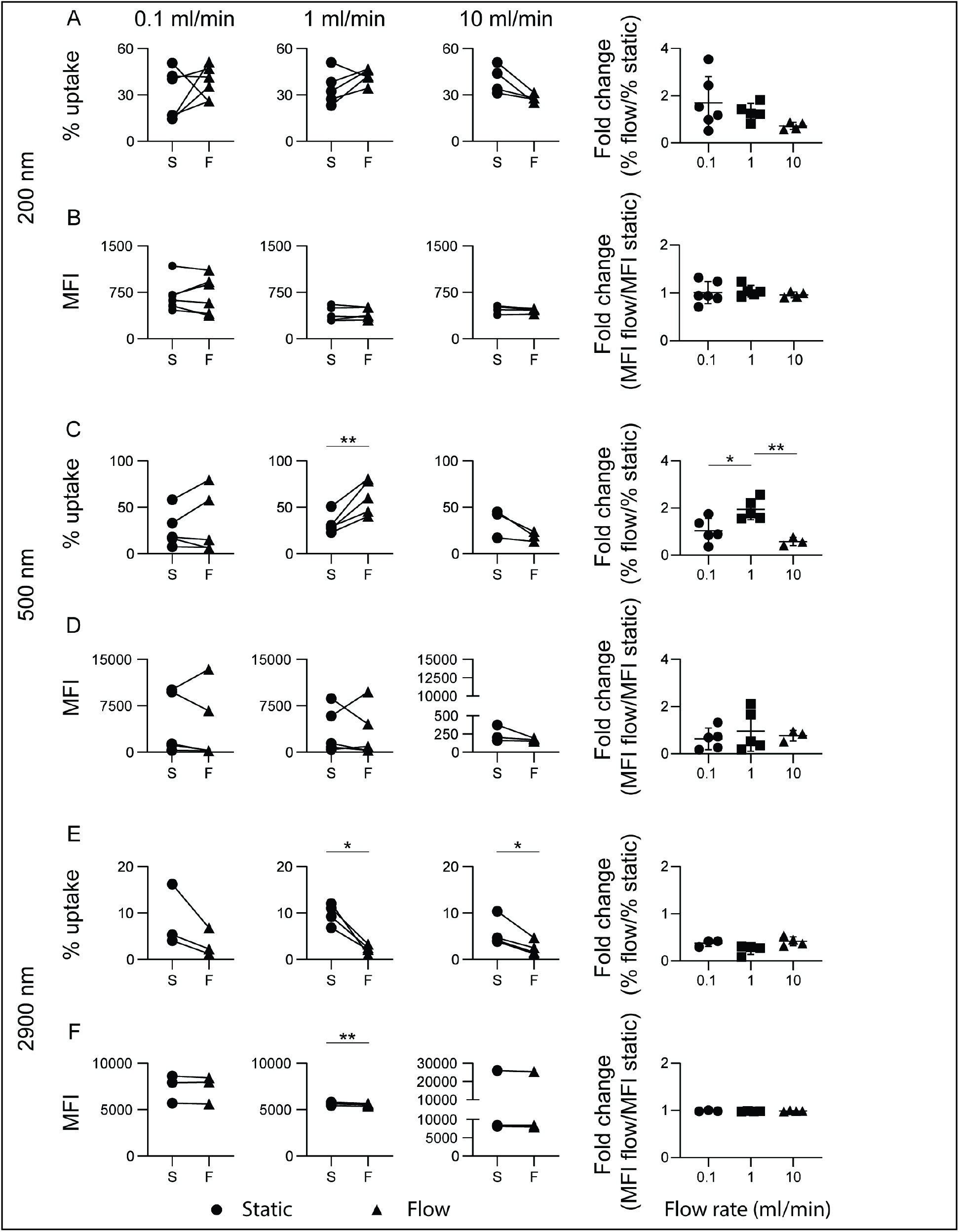
Phagocytosis by the adherent macrophage cell-line (RAW 264.7) at various flow rates. Uptake of 200 nm (**A and B**) or 500 nm (**C and D**) or 2900 nm (**E and F**) particles plotted as either percentage uptake or particles per cell measured as median fluorescence intensity (MFI). Cell to particle ratio of 1:3000 (200 nm) or 1:50 (500 nm) or 1:1 (2900 nm) was used. Data represent results of n≥3 independent experiments. A paired Student’s ‘t’ test was performed to determine statistical differences between two groups and a one-way ANOVA followed by post-hoc Tukey test was performed to determine statistical differences between multiple groups. * = p < 0.05, ** = p < 0.01, and non-significant when no notation.

Further, we observed that under flow conditions, the percentage of cells taking up larger 2900 nm (2.9 µm) sized particles reduced significantly compared to static conditions but was not different across flow rates (Figure 1E). Nevertheless, the small number of cells that did take up 2900 nm particles under flow conditions could phagocytose a similar number of particles as static conditions (Figure 1F). Together, these data show that adherent macrophages’ capacity to take up smaller particles, but not larger particles, under flow conditions remains similar (if not greater) to static conditions.

Having observed that at the flow rate of 1 ml/min, uptake is greater under flow conditions for 500 nm particles, we evaluated if the overall uptake or the kinetics of uptake was greater. Upon determining uptake over time, we observed that while the percentage of cells with particles was higher until 2 hours for 1ml/min flow compared to static conditions (and other flow rates), by 4 hours, the percentage uptake between the two conditions was not statistically different (Figure 2A). Additionally, the number of particles per cell (MFI) remains similar (not statistically different) under flow compared to static conditions (Figure 2B). These data suggest that the overall uptake under flow conditions is not greater. Rather the kinetics of particle-cell interaction resulting in uptake may be higher, resulting in faster uptake at the flow rate of 1ml/min.

**Figure 2:**
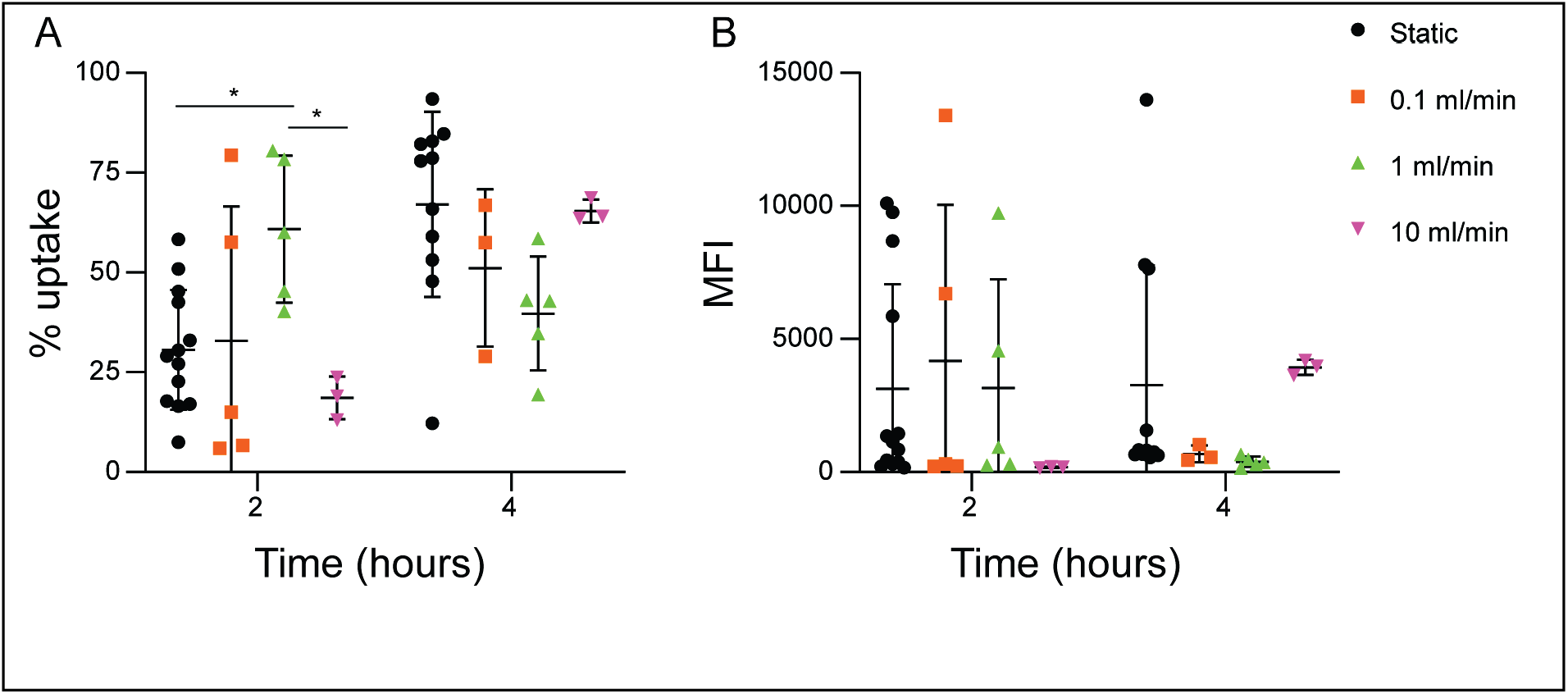
Kinetics of particle uptake. **A** – percentage uptake and **B** – median fluorescence intensity (MFI) denoting particles per cell at two and four hours under static and flow conditions to study rates of uptake. A one-way ANOVA followed by post-hoc Tukey test was performed to determine statistical differences between multiple groups. * = p < 0.05 and non-significant when no notation

### Uptake by a non-adherent cell line

Next, we sought to determine whether cells grown in suspension cultures would also behave similarly to an adherent cell line. We used HL-60 cells^22,23^, a human promyeloblast cell line for these studies, and used a higher ratio of particles to cells as these cells are less phagocytic. We observed that uptake by HL-60 cells also changed based on the size of the particle and flow rate, but the data were different from RAW cell uptake. Among the smaller particles (200 nm and 500 nm), uptake under static and flow conditions were similar at lower flow rates (0.1 and 1 ml/min). However, uptake was significantly lower at the highest flow rate of 10 ml/min (Figure 3A, 3B, 3C and 3D). This data indicates that this cell line cannot engulf particles at flow rates that approximately correspond to flow in large veins or smaller arteries. For the larger-sized particle (2900 nm), the percentage of cells with particles was low under both static and flow conditions (Suppl. Fig. 3), but the differences between the two were minimal (Figure 3E and 3F).

**Figure 3:**
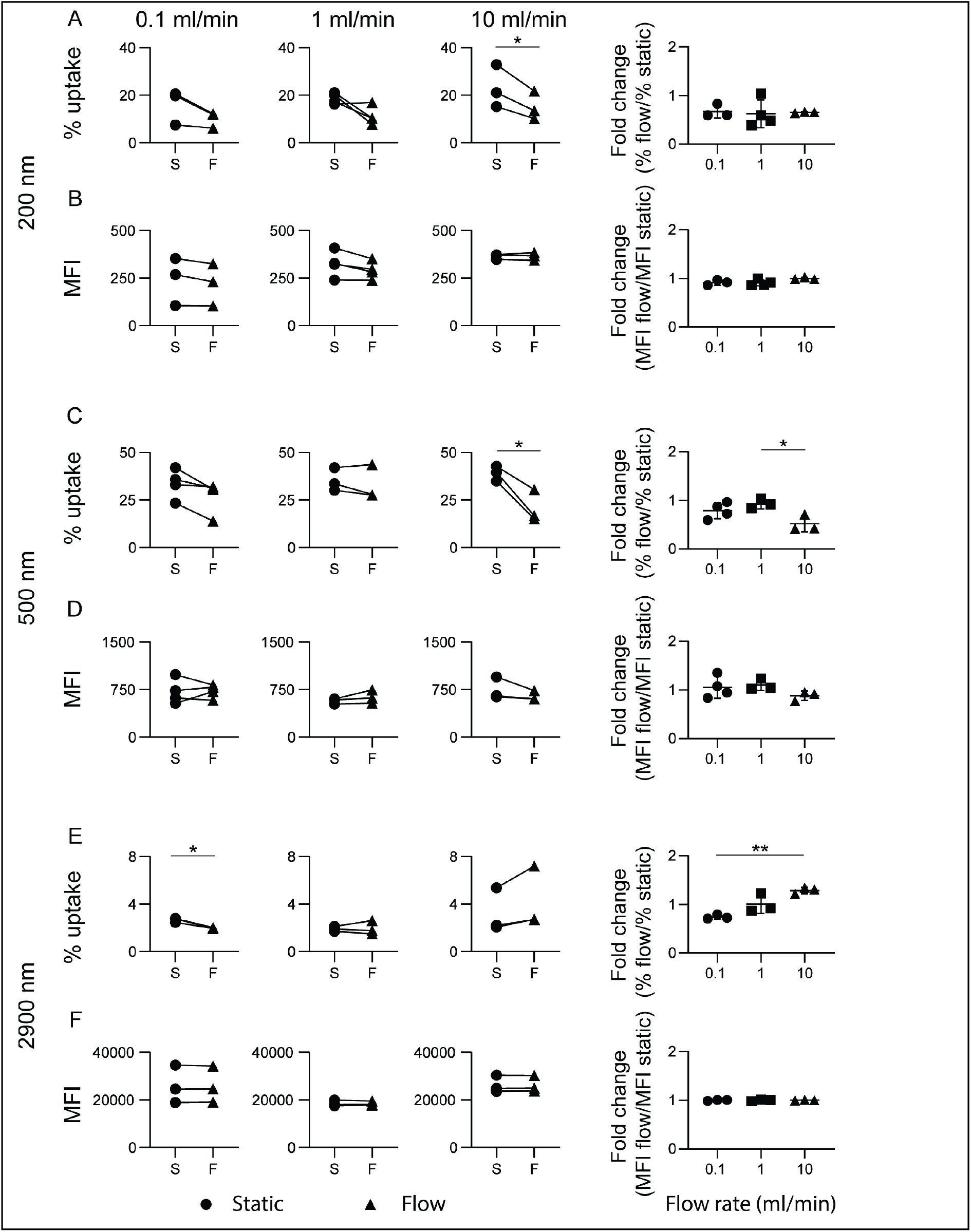
Phagocytosis by the suspension myeloid cell-line (HL-60) at various flow rates. Uptake of 200 nm (**A and B**) or 500 nm (**C and D**) or 2900 nm (**E and F**) PS particles plotted as either percentage uptake or particles per cell measured as median fluorescence intensity (MFI). Cell to particle ratio of 1:9000 (200 nm) or 1:200 (500 nm) or 1:4 (2900 nm) was used. Data represent results of n≥3 independent experiments. A paired Student’s ‘t’ test was performed to determine statistical differences between two groups and a one-way ANOVA followed by post-hoc Tukey test was performed to determine statistical differences between multiple groups. * = p < 0.05, ** = p < 0.01, and non-significant when no notation.

### Uptake by primary cells

The cell line data suggest that particle internalization is possible under flow conditions, but cell lines may not entirely mimic primary cells^24^. Hence, we sought to determine if the observations held for cells isolated from human peripheral venous blood. For these studies, particles of one size (500 nm) were used. Specific immune cells were purified using density gradient centrifugation, and isolated cell populations were subjected to phagocytosis studies under static and flow conditions. We observed that neutrophils took up particles slightly, but not significantly, better under flow than static conditions (Figure 4A, Suppl. Fig. 4). A similar trend was observed for the number of particles per cell (Figure 4B). On the other hand, peripheral blood mononuclear cells (PBMCs) showed a significantly increased percentage of cells with particles at 1 and 10 ml/min compared to static conditions (Figure 4C, Suppl. Fig. 4). Additionally, these cells also took up more particles per cell at the highest flow rate of 10 ml/min (Figure 4D). Nevertheless, the differences were not significant when comparing across flow rates (Figure 4C and 4D).

**Figure 4:**
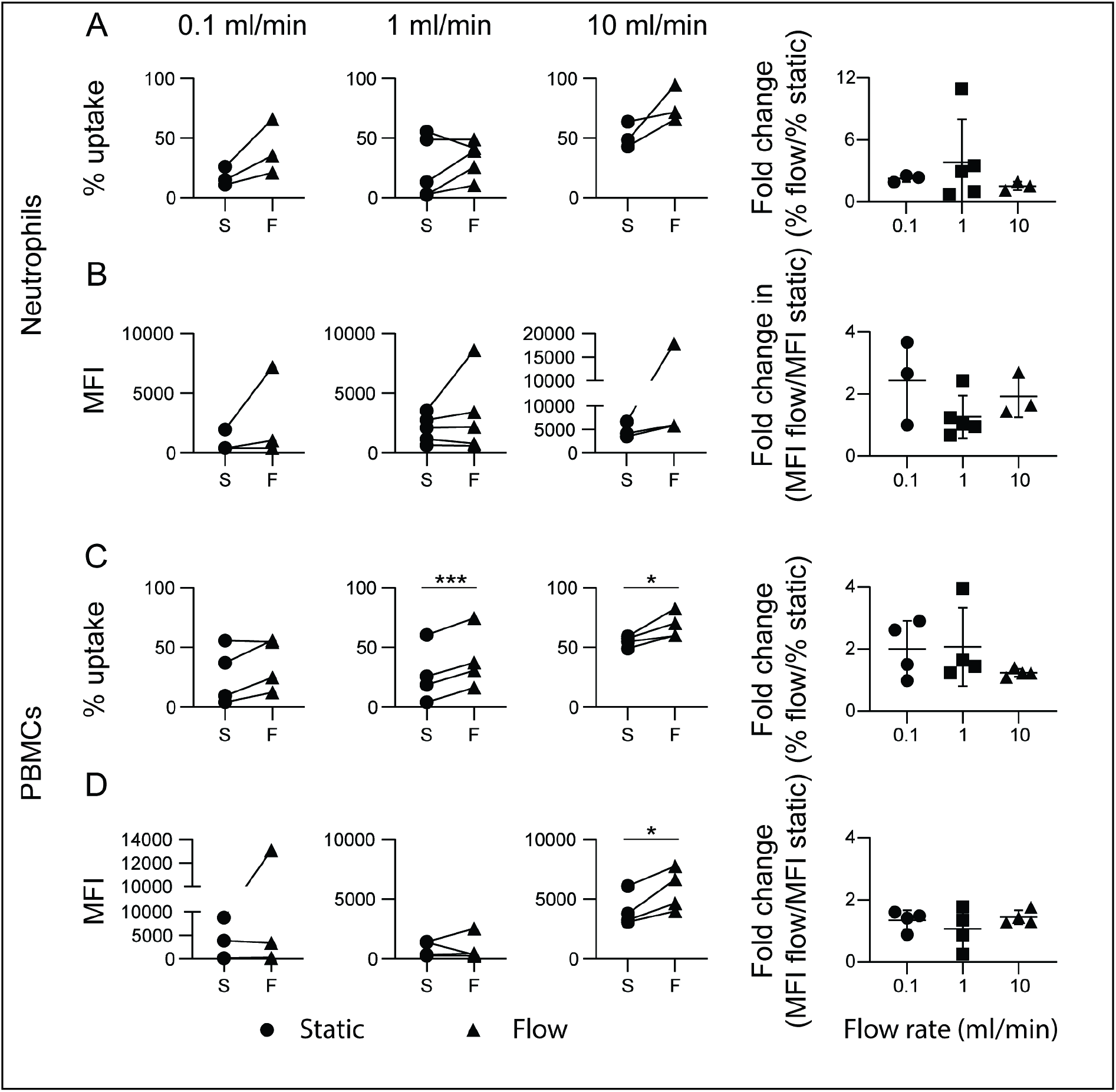
Phagocytosis by human neutrophils and PBMCs. Uptake of 500 nm particles by neutrophils (**A and B**) or PBMCs (**C and D**) isolated from peripheral venous blood draws, plotted as either percentage uptake or particles per cell measured as median fluorescence intensity (MFI). Cell to particle ratio of 1:200 was used. Data represent results of n≥3 independent experiments. A paired Student’s ‘t’ test was performed to determine statistical differences between two groups and a one-way ANOVA followed by post-hoc Tukey test was performed to determine statistical differences between multiple groups. * = p < 0.05, *** = p < 0.001, and non-significant when no notation.

### Whole blood experiments

While the data from purified primary cells suggests that uptake does occur under flow conditions, whole blood is a crowded environment where red blood cells outnumber the phagocytic immune cells by over a thousand to one. Such crowding affects the flow of cells and particulates^25^, which in turn might affect phagocytosis. Particles were added to whole blood, and uptake was measured under both flow and static conditions to determine if uptake occurs under crowded conditions. After co-culture for two hours, cells were stained with antibodies to identify neutrophils and monocytes (Suppl. Fig. 5), and the uptake was measured. Interestingly, neutrophils from whole blood were better at taking up particles under flow conditions compared to static conditions at 0.1 ml/min (significantly better) and 1 ml/min (trending higher but no significant difference) flow rates (Figure 5A). At the higher flow rate of 10 ml/min, this difference was no longer observed, with neutrophils taking up particles equally well under static and flow conditions (Figure 5A). Similar to the observations in most cell line and purified primary cell studies, the total number of particles per cell did not appreciably change at any flow rate compared to static conditions (Figure 5B). Another observation from these studies was that the overall uptake of purified neutrophil populations was higher than neutrophils in whole blood, under both flow and static conditions. Such lowering could be due to a reduced chance of a neutrophil encountering a particulate^24^.

**Figure 5:**
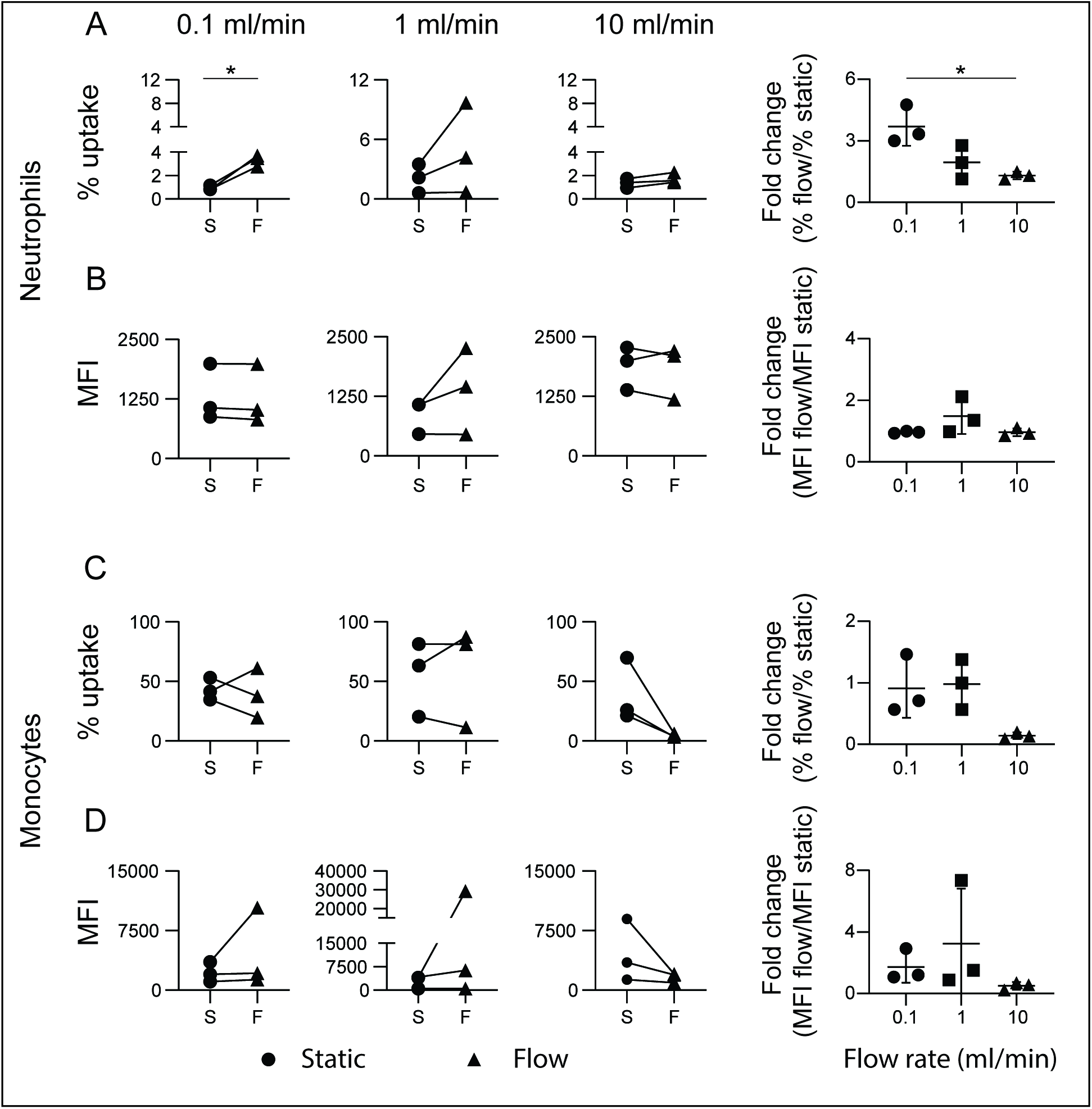
Phagocytosis by immune cells in whole blood. Uptake of 500 nm particles by neutrophils (**A and B**) or monocytes (**C and D**) present in whole blood, plotted as either percentage uptake or particles per cell measured as median fluorescence intensity (MFI). Cell to particle ratio of 1:200 was used according to combined theoretical neutrophil and PBMC counts in the blood (7×10^6^ cells/ml). Data represent results of n=3 independent experiments. A paired Student’s ‘t’ test was performed to determine statistical differences between two groups and a one-way ANOVA followed by post-hoc Tukey test was performed to determine statistical differences between multiple groups. * = p < 0.05 and non-significant when no notation.

In comparison, monocytes from whole blood showed similar uptake at the lower flow rates of 0.1 and 1 ml/min and static conditions (Figure 5C). However, they did show a propensity to have lower uptake at the higher flow rate of 10 ml/min compared to static conditions, but these differences were not statistically different in our studies with a limited number of samples (Figure 5C). This trend continued for the number of particles per cell, with similar numbers observed at 0.1 and 1 ml/min but lower at 10 ml/min flow rate (Figure 5D).

## Discussion

The uptake of particles via one of the endocytic processes occurs in a series of steps^26,27^. The first involves immune cells coming in contact with the particle they intend to take up, which has been thought to occur when the cell crawls towards the particulate substance^28^. The next step is the formation of membrane extensions that engulf the target^29^. And the final steps are the internalization of the particle in an endosomal vesicle. The process of cell crawling and extending its membrane around a particle is more likely when there is minimal to no flow involved. Additionally, it has been suggested that these same cellular processes may also occur when the cell is stationary while the particle is flowing, with the particle’s size and shape influencing the frequency of interactions^30^. However, it remains unclear if the particulate uptake processes mentioned above^28,29^ apply to the condition where both cells and particles are flowing.

Our data shows that uptake does occur when both cells and particles are flowing. How this might happen in the light of the established models of endocytosis is to be determined. We speculate that flow conditions might increase the number of random interactions (collisions) between cells and particles. Preliminary evidence for this statement comes from our data on reduced uptake by neutrophils and monocytes under crowded conditions of whole blood (where productive collision events will be lower) than uptake levels of purified populations of the same cells.

Interactions form only the first step of the uptake process. In the next stage of engulfment, cells must extend their membrane around the particle. This step involves cytoskeletal rearrangements^31^, and whether such changes occur under flow conditions needs to be explored further. Our data of equivalent or higher uptake of the smaller (200 and 500 nm) sized particles under flow conditions suggest that cytoskeletal rearrangement might occur in these conditions. Similarly, lower uptake of the larger (2900 nm) particles might mean that even if such cytoskeletal reorganization occurs, they are likely limited and cannot engulf large particles.

Finally, the data presented here is for particles made of one material (polystyrene). While the phenomena of uptake, when both cells and particles are flowing, may apply to hard particles made of similar polymers and possibly even metallic materials, it remains to be seen if it would apply to softer particles such as liposomes. The particle shape also influences uptake^32^, and we study particles of one shape (spherical). Future studies will need to examine how particle shape might alter their flow dynamics and affect uptake under fluid flow conditions.

In conclusion, we show that immune cells are capable of taking up particles under flow conditions. The uptake capacity is either similar or greater while flowing than static conditions for sub-micrometer-sized particles but lower for larger micro-sized particles. Neutrophils and monocytes are also capable of taking up particles under the crowded conditions of whole blood. These studies reveal that circulating immune cells do have the capacity to take up particulates injected intravenously, which has potential implications in our design of delivery vehicles that either seek to avoid or target these cells.

## Acknowledgments

We thank G. K. Ananthasuresh for providing the peristaltic pump system. This work was supported by the Dr. Vijaya and Rajagaopal Rao funding for Biomedical Engineering research at the Centre for BioSystems Science and Engineering and the R. I. Mazumdar young investigator position (to SJ). This work was also supported in part by the DBT-Wellcome India Alliance Intermediate Fellowship number IA/I/19/1/504265 (to SJ). MS and PS are in receipt of the Kishore Vajgyanik Protsahan Yojana and CSIR senior research fellowship, respectively.

