## Supplementary for "Phagocytic Uptake of Particles by Immune cells Under Flow Conditions"

Supplementary Figure 1

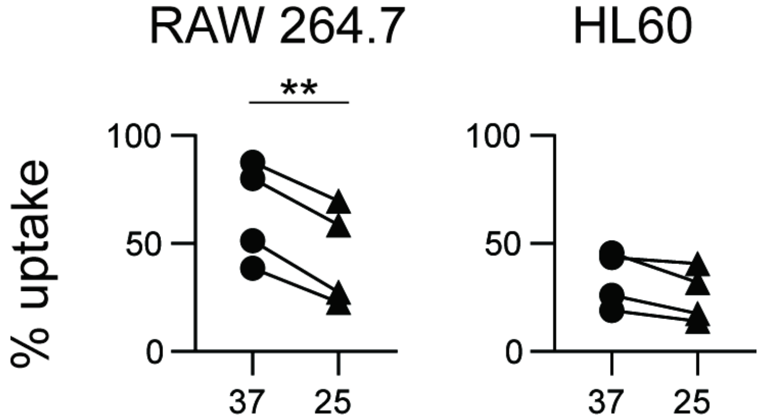


**Supplementary Figure 1:** Comparison of phagocytosis of 500 nm PS particles by RAW 264.7 or HL60 cell lines at two different temperatures (37 degree and 25 degree) under static conditions. A paired Student's 't' test was performed to determine statistical differences between two groups. * = p <0.05, ** = p<0.01 and non-significant when no notation

Supplementary Figure 2

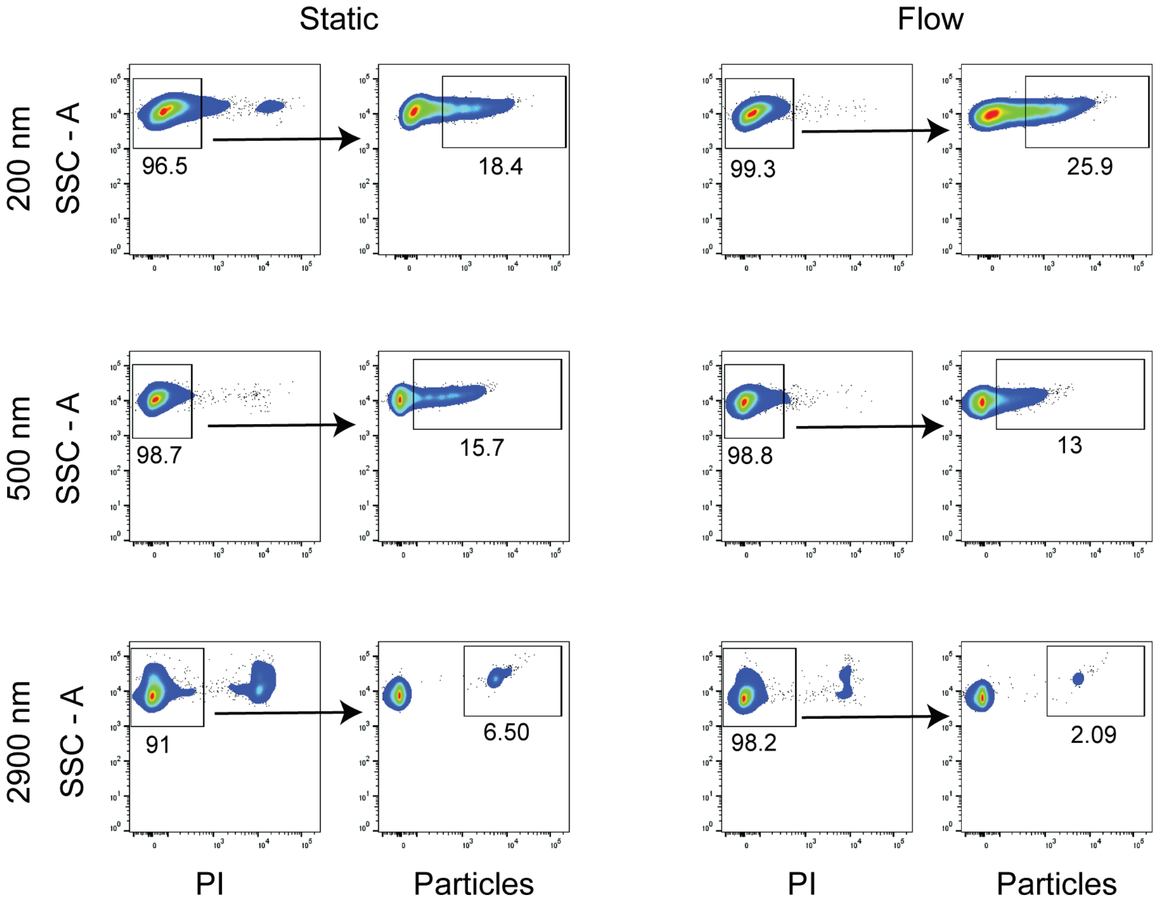


**Supplementary Figure 2:** Flow cytometry density plots (of RAW 264.7 cells with particles) showing gating on live cells (left) followed by gating on the population of cells with particles (right). Single color controls were used to determine gate location. PI denotes propidium iodide, and particles denotes the particle fluorescence

Supplementary Figure 3

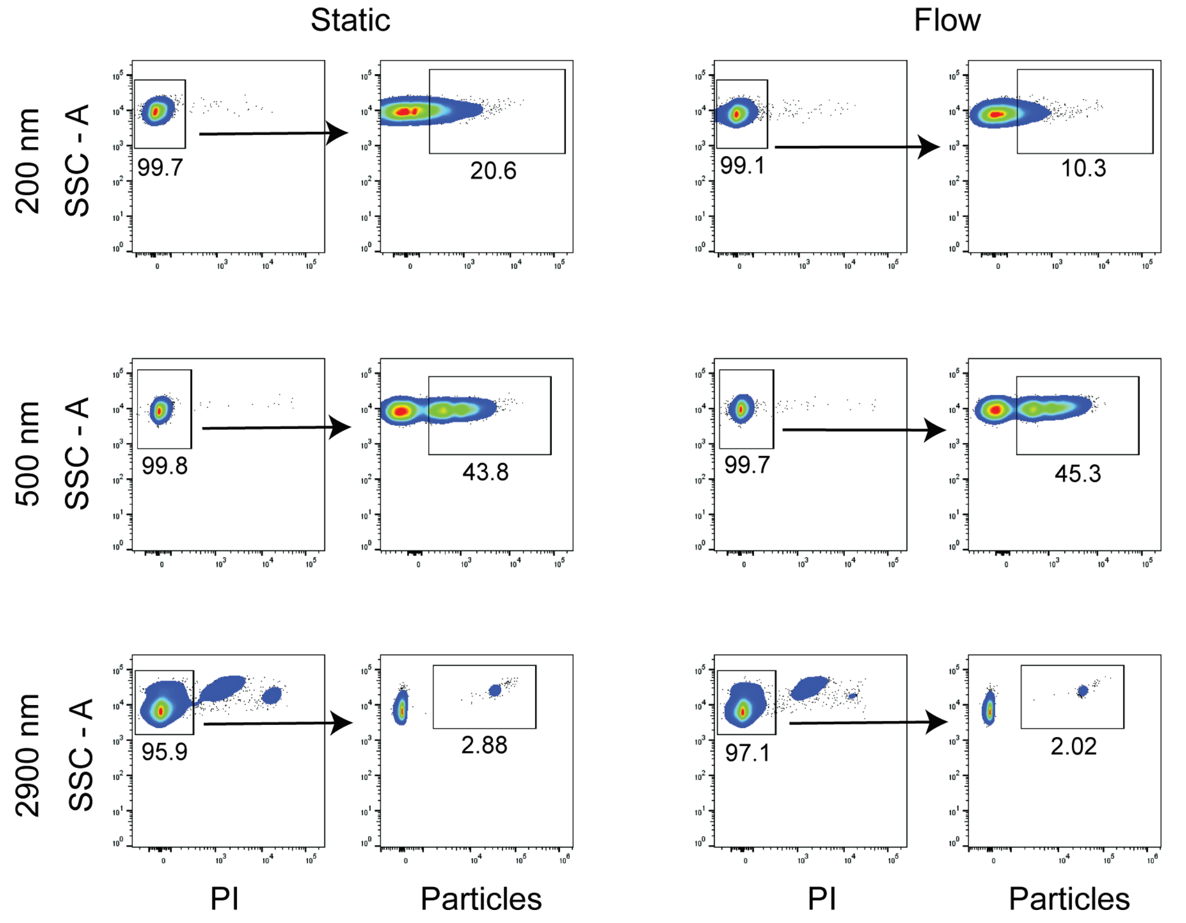


**Supplementary Figure 3:** Flow cytometry density plots (of HL60 cells with particles) showing gating on live cells (left) followed by gating on the population of cells with particles (right). Single color controls were used to determine gate location. PI denotes propidium iodide, and particles denotes the particle fluorescence

Supplementary Figure 4

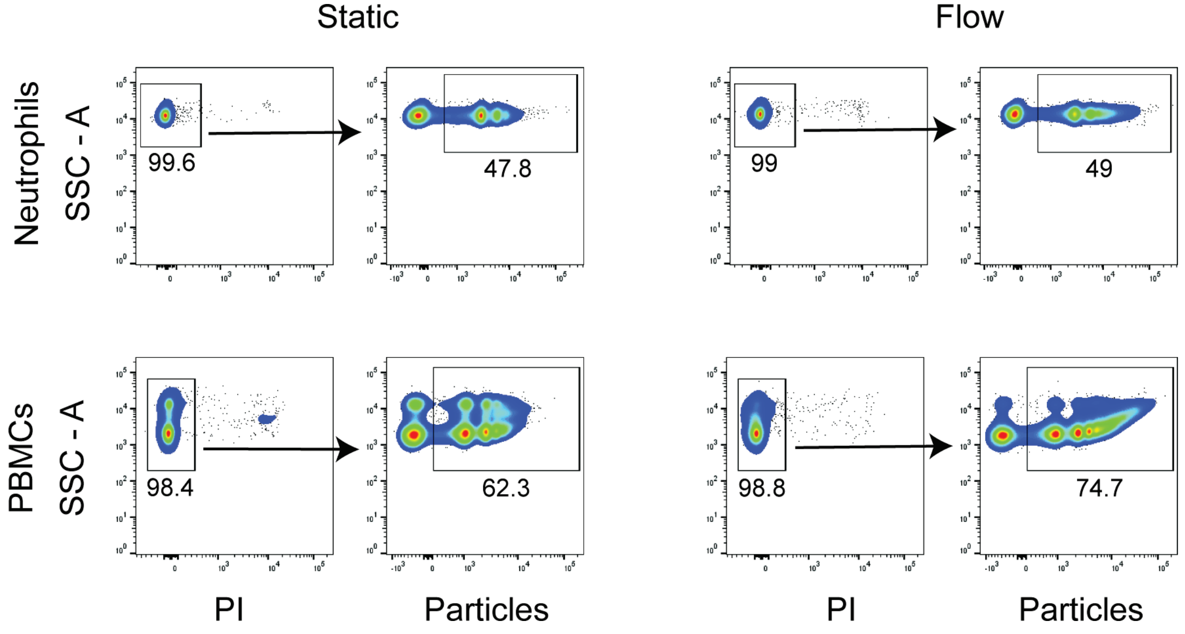


**Supplementary Figure 4:** Flow cytometry density plots (of isolated primary cells with particles) showing gating on live cells (left) followed by gating on the population of cells with particles (right). Single color controls were used to determine gate location. PI denotes propidium iodide, and particles denotes the particle fluorescence

Supplementary Figure 5

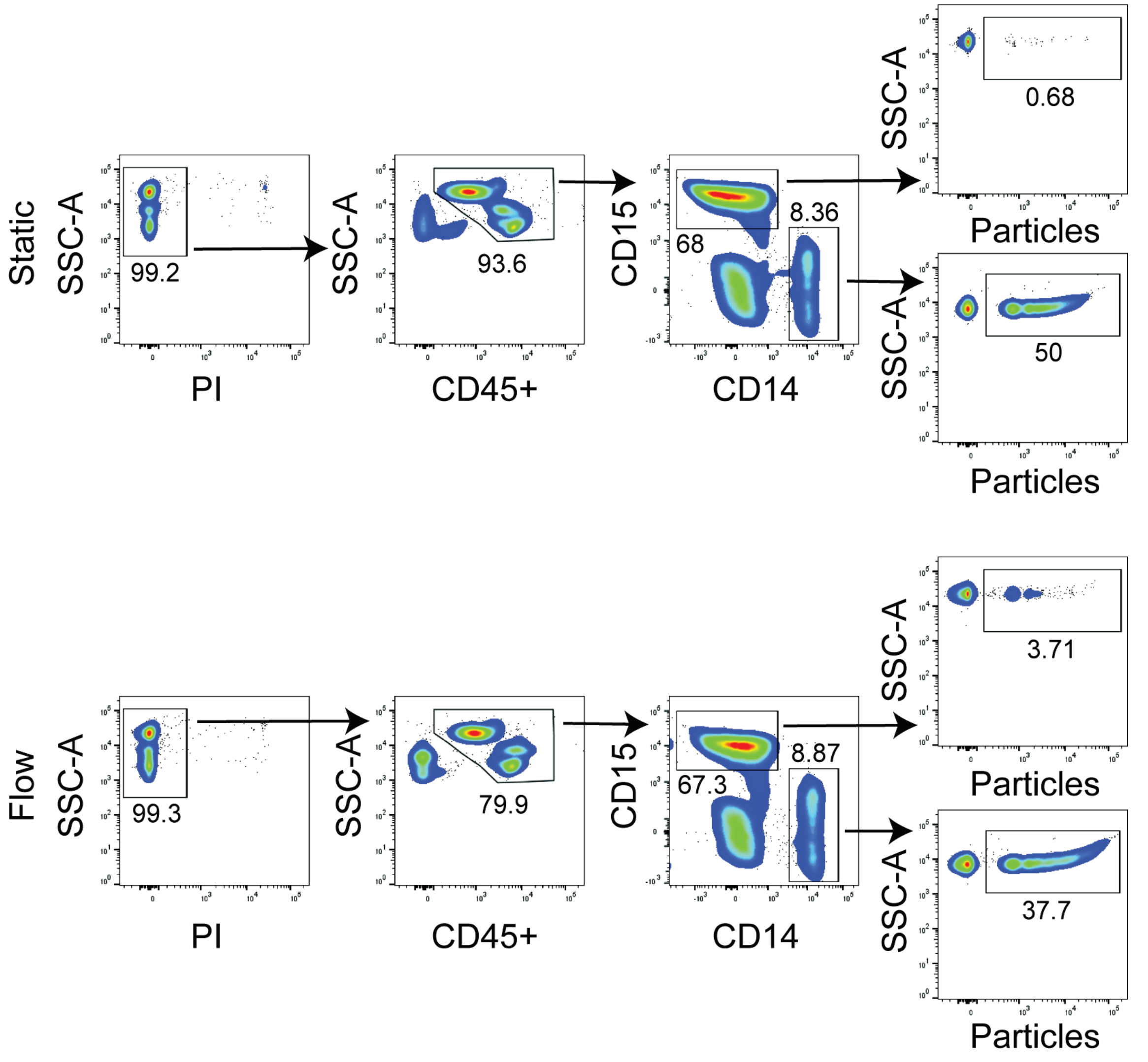


**Supplementary Figure 5:** Flow cytometry density plots (of cells from whole blood) showing gating strategy to determine immune cells (CD45+), and monocytes (CD14+ CD15-) and neutrophils (CD15+ CD14-). Uptake was measured among the neutrophils and monocytes under both static and flow conditions. Single color controls were used to determine gate location. PI denotes propidium iodide, and particles denotes the particle fluorescence
